# Automated Identification of Germline *de novo* Mutations in Family Trios: A Consensus-Based Informatic Approach

**DOI:** 10.1101/2024.03.08.584100

**Authors:** Mariya Shadrina, Özem Kalay, Sinem Demirkaya-Budak, Charles A. LeDuc, Wendy K. Chung, Deniz Turgut, Gungor Budak, Elif Arslan, Vladimir Semenyuk, Brandi Davis-Dusenbery, Christine E. Seidman, H. Joseph Yost, Amit Jain, Bruce D. Gelb

**Affiliations:** Mindich Child Health and Development Institute and the Department of Genetics and Genomic Sciences, Icahn School of Medicine, New York, NY, USA; Velsera Inc, 529 Main St, Suite 6610, Charlestown, MA, USA; Department of Pediatrics, Columbia University, New York, NY, USA; Department of Pediatrics, Boston Children’s Hospital, Harvard Medical School, Boston, MA, USA; Division of Cardiovascular Medicine, Brigham and Women’s Hospital, Harvard Medical School, Boston, MA, USA; Howard Hughes Medical Institute, Chevy Chase, MD, USA; Molecular Medicine Program, University of Utah, Salt Lake City, UT, USA; Department of Pediatrics, Icahn School of Medicine, New York, NY, USA

## Abstract

Accurate identification of germline *de novo* variants (DNVs) remains a challenging problem despite rapid advances in sequencing technologies as well as methods for the analysis of the data they generate, with putative solutions often involving *ad hoc* filters and visual inspection of identified variants. Here, we present a purely informatic method for the identification of DNVs by analyzing short-read genome sequencing data from proband-parent trios. Our method evaluates variant calls generated by three genome sequence analysis pipelines utilizing different algorithms—GATK HaplotypeCaller, DeepTrio and Velsera GRAF—exploring the assumption that a requirement of consensus can serve as an effective filter for high-quality DNVs. We assessed the efficacy of our method by testing DNVs identified using a previously established, highly accurate classification procedure that partially relied on manual inspection and used Sanger sequencing to validate a DNV subset comprising less confident calls. The results show that our method is highly precise and that applying a force-calling procedure to putative variants further removes false-positive calls, increasing precision of the workflow to 99.6%. Our method also identified novel DNVs, 87% of which were validated, indicating it offers a higher recall rate without compromising accuracy. We have implemented this method as an automated bioinformatics workflow suitable for large-scale analyses without need for manual intervention.

## INTRODUCTION

Germline *de novo* mutations (DNV) play a crucial role in evolution, introducing new genetic variation. At the same time, DNVs underlie a wide range of genetic diseases, increasing the interest in studying the frequency and characteristics of sporadic mutations in human genomes (Acuna-Hidalgo et al., 2016; Deciphering Developmental Disorders Study, 2017; Goldmann et al., 2019). With the recent availability of genome sequencing (GS), genetic studies of trios consisting of an affected proband and unaffected parents provide a direct method for the large-scale detection of DNVs (Nicolas & Veltman, 2019; Richter et al., 2020). Although the genome sequence of an individual can differ at 4-5 million positions compared to the human reference genome (The 1000 Genomes Project Consortium et al., 2015), the vast majority of the observed genetic variation is inherited. The germline *de novo* mutation rate for single nucleotide variants (SNVs) in human genomes is estimated as 1.0–1.8 × 10^−8^ per nucleotide per generation, which manifests as 44 to 82 *de novo* SNVs for an individual (including one to two variants in coding regions) and is dependent upon parental ages, predominantly paternal age (Acuna-Hidalgo et al., 2016; Goldmann et al., 2019). In addition to SNVs, only three to nine small *de novo* insertions/deletions (indels), which are typically shorter than 50 bp, are expected per human genome. As a result, the prior odds of a variant observed only in the proband genome being a DNV remains modest. Outnumbered by inherited variants, detection of DNVs is a non-trivial task, resulting in many false-positive variant calls, especially in regions of low coverage or with high levels of noise.

In our previous work, we studied 763 probands with congenital heart disease (CHD) and their unaffected parents with trio GS (Richter et al., 2020). We identified 71 *de novo* SNVs and five *de novo* indels per CHD proband on average, corresponding to expected rates of true *de novo* SNVs and indels (around 98% and 94% respectively, based on PCR-based Sanger sequencing). However, accurate detection of DNVs with high precision and sensitivity was achieved using a sophisticated workflow that included manual inspection of ambiguous variants. This limits the scalability of that method for studies of larger cohorts with trio GS, which are becoming increasingly commonplace as costs have decreased. Here, we report the development of a fully automated trio GS workflow implementing three independent pipelines, Broad Institute’s Best Practices Pipeline for Germline Short Variant Discovery (GATK4) (DePristo et al., 2011), Velsera GRAF Germline Variant Detection Workflow (GRAF) (Rakocevic et al., 2019), and BWA- DeepTrio (Kolesnikov et al., 2021), to accurately call DNVs.

## METHODS

GS data from 10 parent offspring trios from the Pediatric Cardiac Genetics Consortium (PCGC) database were analyzed. Each trio consisted of an individual with CHD and their healthy parents. The approach for DNA extraction and GS has been previously described (Richter et al., 2020). In brief, paired-end, short-read genome sequencing was performed with a HiSeq X Ten System (Illumina Inc., San Diego, CA) and achieved average coverage of 35-40x for all samples. To analyze the GS trio data, we ran three analytic pipelines to call *de novo* SNVs and indels: GATK4 and DeepTrio, which rely upon alignment to the single haplotype reference genome assembly using BWA-MEM (Li, 2013), and GRAF, which uses alignments to a pangenome reference. **Figure 1**, **Figure S1** and **Table S1** show all steps performed in the analysis. Human reference genome assembly GRCh38 (Schneider et al., 2017) was used as the basis for variant calls in all pipelines.

**Figure 1.**
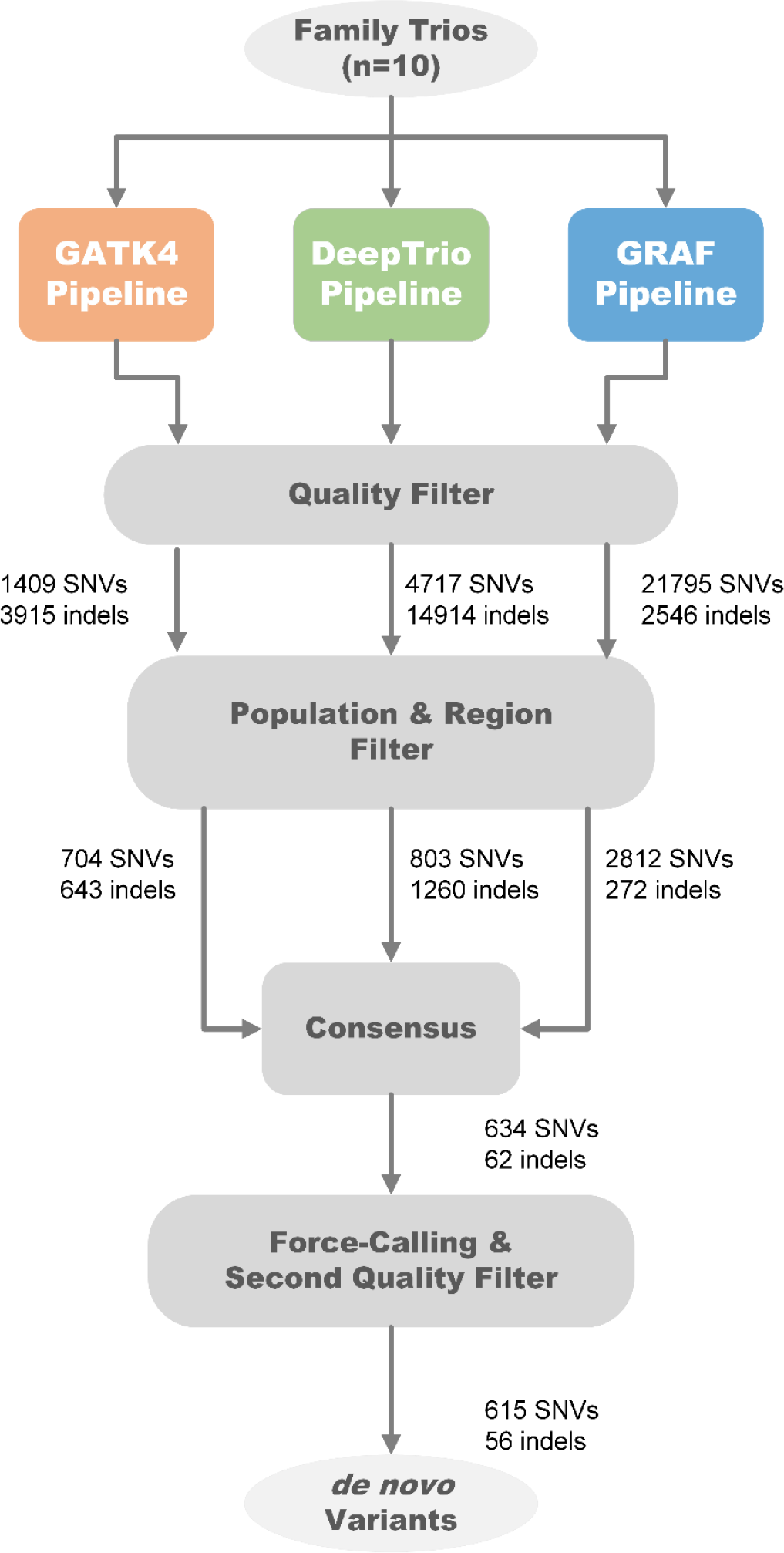
Three pipelines were applied to analysis 10 trios: GATK4, GRAF, DeepTrio. Three independent sets of possible *de novo* variants in probands were found for each family, then filtered with regional and population filters. Only variants found by at least two pipelines were kept. At the last step, the force-calling filter was performed.

*GATK4 pipeline.* The GATK4 pipeline was constructed following the latest version of the Broad Institute’s Best Practices Workflow for germline short variant discovery (**Figure 1**, **Table S1**), a standard approach to small variant calling with linear reference genomes. Paired-end reads were mapped using BWA-MEM followed by variant calling with HaplotypeCaller (Poplin et al., 2017). gVCF files generated for each family member were jointly genotyped using GATK GenotypeGVCFs (van der Auwera & O’Connor, 2020). Variant Quality Score Recalibration (VQSR) and Genotype Refinement steps were applied next. Possible *de novo* calls were annotated with VariantAnnotator.

*DeepTrio pipeline.* DeepTrio is a machine learning-based variant caller that analyzes family trio alignments together (Kolesnikov et al., 2021). It employs deep convolutional neural networks to learn variant context and *de novo* rate from trio data and then utilizes this model to call variants using trio alignments. We used BWA-MEM generated alignments of the family trio as input to DeepTrio variant caller (**Figure 1**, **Table S1**), and the resulting gVCF files for all three family members were jointly genotyped using GLnexus (Yun et al., 2020).

*GRAF pipeline.* Velsera GRAF Germline Workflow utilizes a pangenome reference for incorporating genomic variation in the secondary analysis process, enabling reduced reference bias [Supplementary Materials – Graph Pangenome Reference]. In this work the GRAF pipeline was used with a human pangenome reference incorporating genetic variation posited by large studies of diverse cohorts (Katsnelson, 2010; Mallick et al., 2016; Mills et al., 2006). The paired-end reads from mother, father and the proband were mapped using the GRAF Aligner to the pangenome reference, and the GRAF VariantCaller was used for calling variants (**Figures 1**, **S2**, **S3**). The VCF files for trio members were merged.

### The First QC Step

We applied hard-threshold filtering using the variant annotations from joint VCFs, to eliminate low-quality calls from GATK4 and DeepTrio outputs. For GRAF outputs, we followed a similar hard-thresholding step using annotations from merged VCF and read alignments, after handling representation differences between variants from each family member. [Supplementary Materials – GRAF *de novo* Variant Detection Pipeline]. The details of annotations and thresholds used for each pipeline are in **Tables S2** and **S3**.

The resulting candidate *de novo* variants after the initial filtering steps comprised 1,409 SNVs and 3,915 indels for GATK4, 4,717 SNVs and 14,914 indels for DeepTrio, and 21,795 SNVs and 2,546 indels for GRAF. The number of filtered Mendelian-inconsistent variant calls at this stage exceeds the expected count of DNVs in 10 probands by at least an order of magnitude (Acuna-Hidalgo et al., 2016). Additionally, these variants show enrichment in indels compared to the background distribution due to our filtering method’s aggressive removal of irrelevant SNPs.

### Regional and Population Filters

After the initial filtering steps, candidate variants from the three pipelines were further refined with regional and population filters. The regional filter removed variants located in low-complexity regions, low-mappability regions (Karimzadeh et al., 2018), ENCODE blacklists (The ENCODE Project Consortium, 2012) and segmental-duplication regions (Vollger et al., 2022). The population filter removed all variants with allele frequencies > 0.1% based on the gnomAD exome (v2.1.1) (Karczewski et al., 2020), gnomAD genome (v2.1.1) (Karczewski et al., 2020) and 1000 Genome (Katsnelson, 2010) databases, as variants with high frequency are unlikely to be pathogenic for most Mendelian traits. The final GATK4 candidate DNVs included 704 SNVs and 643 indels; DeepTrio candidate DNVs included 803 SNVs and 1260 indels; and the GRAF candidate DNVs included 2,812 SNVs and 272 indels (**Figure 1**). The union set of DNVs from all three workflows contained 3,120 SNVs and 2,071 indels.

### Consensus Step

Following regional and population filters, we observed that almost all high-confidence DNVs from the previous work (Richter et al., 2020) (Freeze variants) were called by at least two pipelines. Given that our primary focus in this work was to enhance precision in *de novo* calling, we discarded all variants identified by a single method. As a result, 634 of 3,120 SNVs and 62 of 2071 indels were retained (**Figure 1**), a total of 696 putative DNVs across the 10 trios (i.e., 69.6 DNVs/proband).

### False Positive Variants

Visual inspection of the 696 candidate DNVs using the BAM files revealed that 23 variants were highly likely to be inherited or alignment errors (**Table 1, GATK4 + GRAF + DeepTrio**). To improve our informatics-based filtering, we studied the characteristics of these variants to develop additional filtering criteria.

**Table 1.**
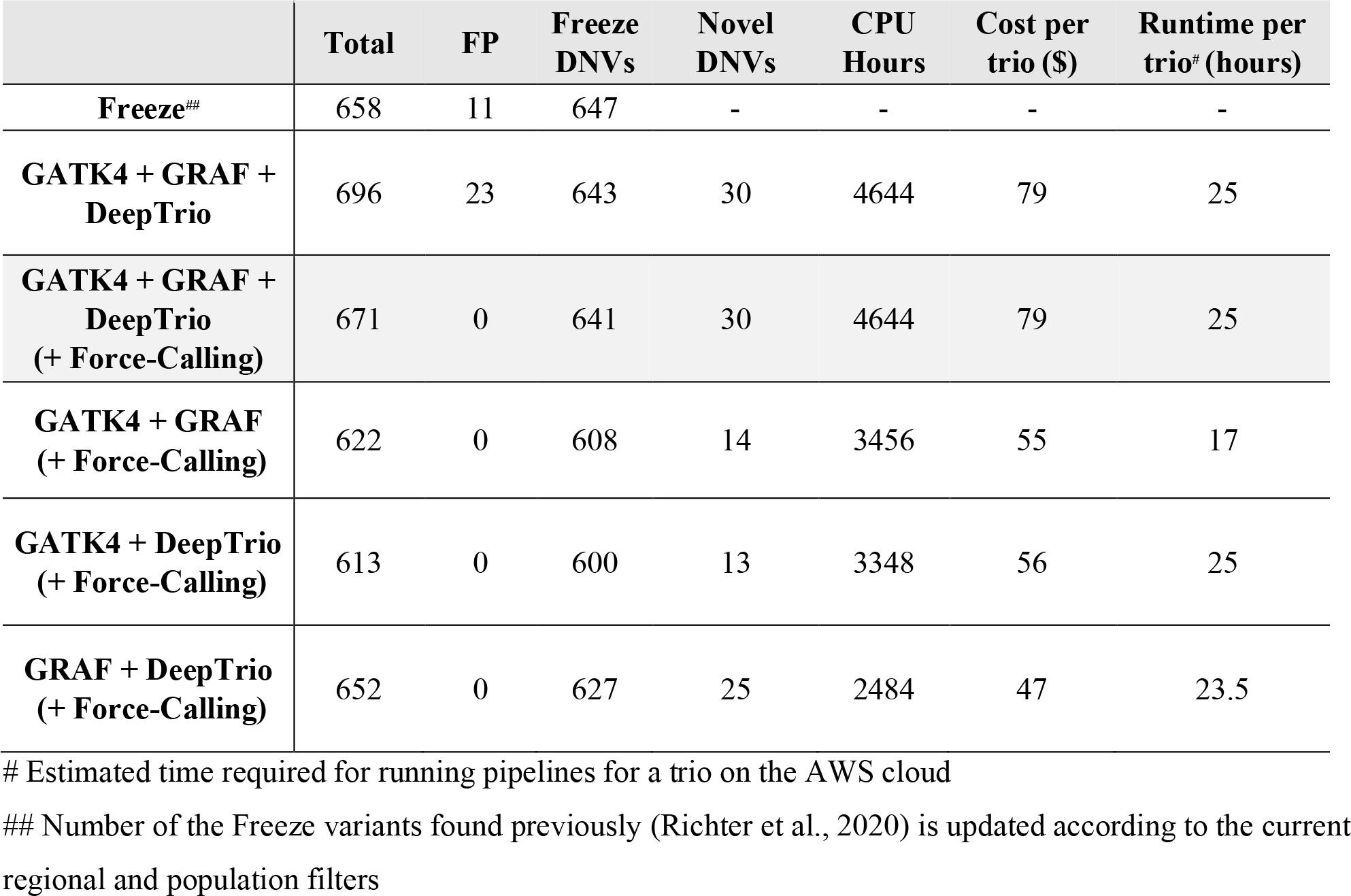
Comparison of combinations of the three pipelines and freeze set.

| | **Total** | **FP** | **Freeze DNVs** | **Novel DNVs** | **CPU Hours** | **Cost per trio (\$)** | **Runtime per trio<sup>#</sup> (hours)** |
| --- | --- | --- | --- | --- | --- | --- | --- |
| **Freeze<sup>##</sup>** | 658 | 11 | 647 | - | - | - | - |
| **GATK4 + GRAF + DeepTrio** | 696 | 23 | 643 | 30 | 4644 | 79 | 25 |
| **GATK4 + GRAF + DeepTrio (+ Force-Calling)** | 671 | 0 | 641 | 30 | 4644 | 79 | 25 |
| **GATK4 + GRAF (+ Force-Calling)** | 622 | 0 | 608 | 14 | 3456 | 55 | 17 |
| **GATK4 + DeepTrio (+ Force-Calling)** | 613 | 0 | 600 | 13 | 3348 | 56 | 25 |
| **GRAF + DeepTrio (+ Force-Calling)** | 652 | 0 | 627 | 25 | 2484 | 47 | 23.5 |

*Alternative alleles in homozygous variants in parents (AAHP filter).* Of the 23 false-positive (FP) variants, nine variants had alternate allele-carrying reads in parents’ pileups even though the variant calls were homozygous reference in the parents (**Figure 2** – A). We considered that many of these variants resulted from alignment errors, where alternative alleles having lower alignment score than the corresponding reference alleles were partially missed by an aligner. Therefore, the discovery of the alternate allele in the proband genotype due to misalignment increases the likelihood of parent genotype containing the alternate allele. However, a single read in a parent alignment showing an alternate SNV coinciding with a *de novo* mutation in the proband may result from technical errors commonly associated with the sequencing process. Considering the latter cases, we applied a threshold for alternative allele carrying reads (AAC) <= 1 for SNVs and 0 for indels in the parent samples. Although these reads could also indicate low-level parental mosaicism, we retained them because that would still be consistent with high-impact variants of clinical significance (Cook et al., 2021).

**Figure 2.**
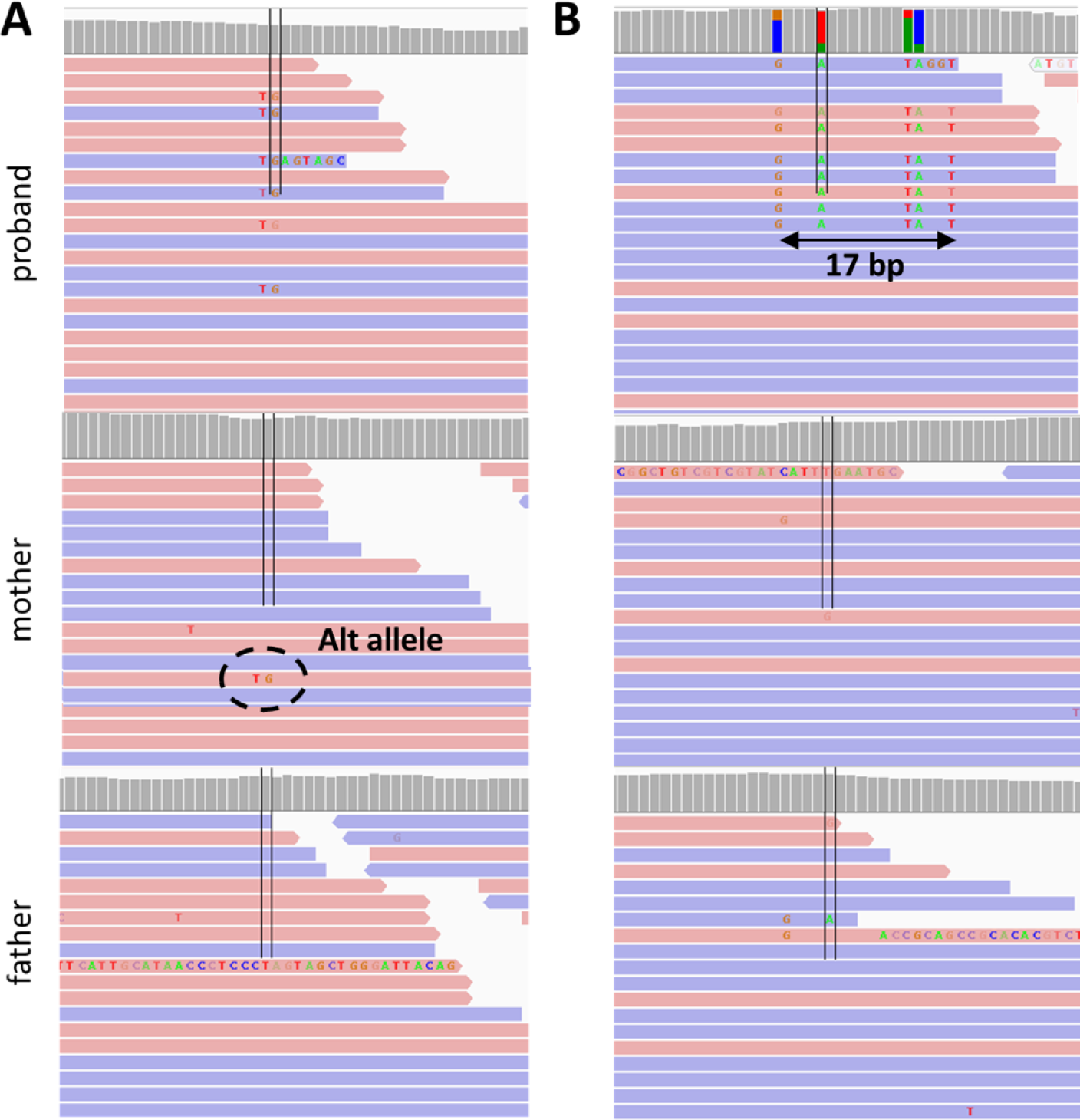
Main indicators of false-positive variants, which are removed after force-calling and second QC. **A.** Reads containing alternative alleles in parental read alignments. **B.** Accumulation of *de novo* SNVs in a relatively restricted genomic area.

*Proband haplotype variants (PH filter).* We observed that 13 variants are in *de novo* clusters, a notable concentration of *de novo* SNVs within a relatively confined genomic region. (**Figure 2B**). We considered that such occurrences might stem from alignment errors, as a substantial number of multiple proximate *de novo* events are unlikely. On the other hand, we did not want to discard the possibility that some neighboring *de novo* mutations might be synchronous. Considering these factors, we implemented a filtering process to exclude DNV clusters that extend beyond five base pairs.

### Force-calling and the Second QC Step

By default, the HaplotypeCaller performs variant calling in regions that have evidence for genomic variation, termed active regions. After determining that the region is active, HaplotypeCaller reassembles the reads, creates a sequence for the region and determines the allele. HaplotypeCaller also allows for reassembling the reads at a given region even if the region is not active (--force-call runtime parameter).

We used this functionality to conduct a more thorough examination of candidate *de* novo variants identified during the consensus step and to search for variant evidence in parents. In this work, we will refer to this step as ‘force-calling’.

Following the application of the force-calling step, we implemented a second round of quality filtering and applied AAHP and PH filters. **Table S2** shows additional QC filters applied to the candidate variants after the force-calling step. After the second QC filtering, the total number of *de novo* variants was reduced to 671 variants (**Table 1, GATK4 + GRAF + DeepTrio + Force-Calling**). A subset of DNVs classified as TP and FP underwent validation in the proband and both parents with Sanger sequencing of amplicons after PCR amplification using primers designed within 100-400 bp of the variant (Zaidi et al., 2013).

## RESULTS

### Comparison of Pipeline Combinations

Most of the final DNV set were found by all three methods: 560 SNVs and 48 indels (**Figure 3**). However, the GRAF pipeline added 14 and 44 DNVs when overlapped with GATK4 and DeepTrio, respectively. In contrast, the consensus between GATK4 and DeepTrio added only five DNVs.

**Figure 3.**
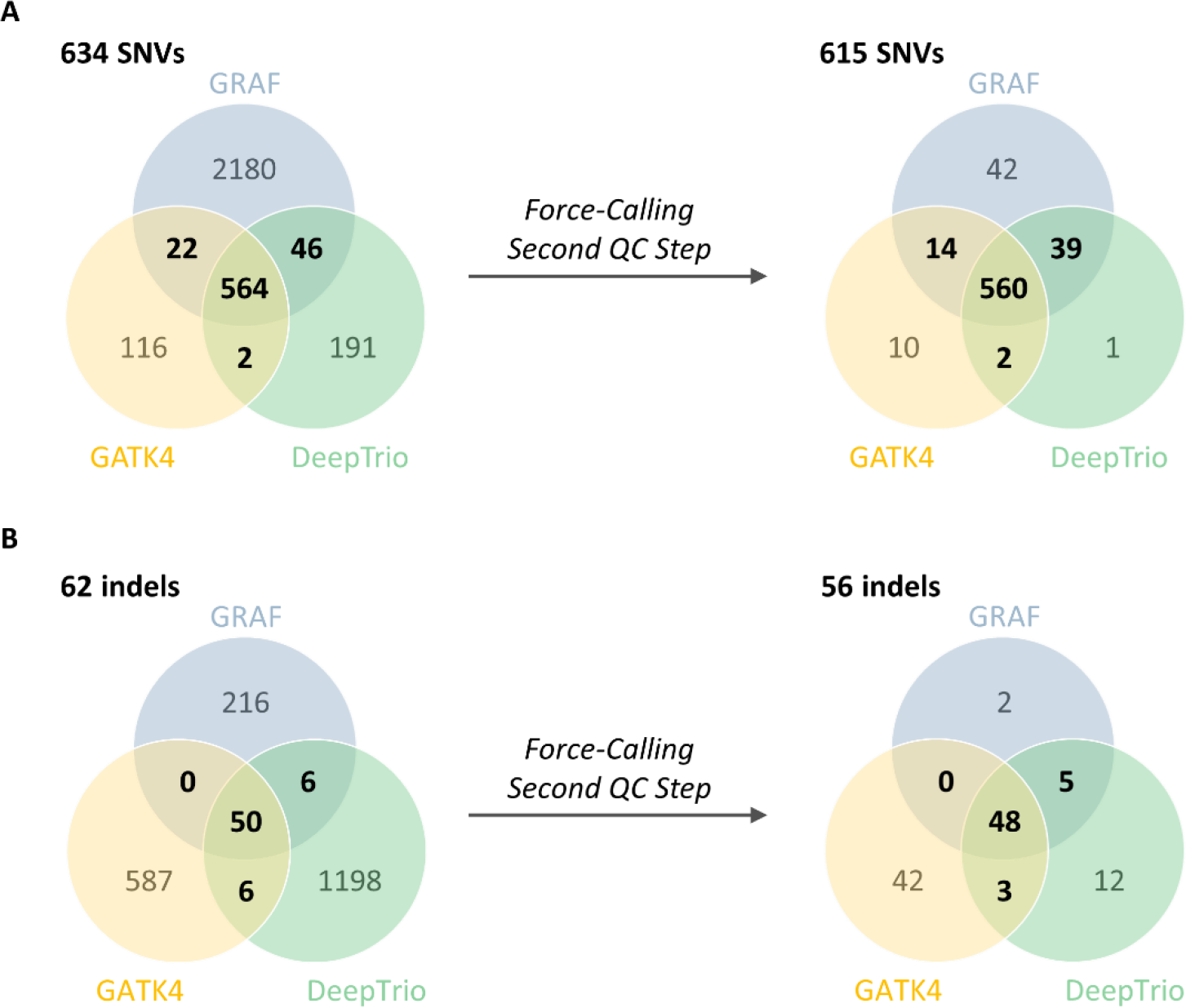
Distribution of the *de novo* candidates before and after the force-calling step between the three pipelines: **A**. SNVs and **B**. indels. Only variants found by at least two methods were included in the analysis.

The consensus workflow combining the results from the three orthogonal variant calling methods showed the best results with the largest total number of DNVs (671 variants), though it was also the most computationally expensive to run (**Table 1**). The pipeline with all three methods on average required 4,644 CPU hours to run on a computer with 72 Intel Xenon CPUs, costing $79 per trio on the AWS cloud platform. Using GRAF and DeepTrio together identified 652 DNV variants (missing 2.8% of variants from the three-method set) but only required 2,484 CPU hours ($47) per trio, while the GATK4 and DeepTrio combination was the least effective, identifying only 613 DNVs (missing 8.6% of DNVs found by the three-method option) while requiring 3,348 CPU hours ($56). Combining GATK4 and GRAF revealed 622 DNVs (missing 7.3% of the three-method set) and taking 3,456 CPU hours ($55) per trio. **Figure S1** and **S4** show the cost and CPU hours distribution over different pipelines.

### Comparison with the Freeze Set

The previous PCGC study called DNVs with a combination of GATK, FreeBayes and a convolutional neural network trained on manually curated IGV plots (Richter et al., 2020).That pipeline found 752 variants for the 10 trios (freeze set), with a true DNV call rate of > 95% based on Sanger sequencing confirmation of a modest number of DNV calls. We reassessed those freeze set variants by applying regional and population filters, which narrowed the DNV set to 658 variants. Eleven variants did not pass visual verification of the BAM files in IGV and were removed as FPs. The remaining 647 Freeze variants were considered as true DNVs and used for comparison (**Table 1**).

Combining all three pipelines (GATK4, DeepTrio, and GRAF) detected 641 of the 647 Freeze variants (**Table 1, GATK4 + GRAF + DeepTrio + Force-Calling**). Two freeze set *de novo* SNVs were called only by the GRAF pipeline and were excluded in the consensus step, and one freeze set *de novo* indel was called only by the GATK4 method. Another freeze set *de novo* indel was missed by all three pipelines. Two freeze set DNVs were additionally filtered out by the PH filter, as the length of the cluster was estimated as 6 bp. In summary, combining all three pipelines and using the force-calling step identified 641 Freeze DNVs (99.1%) and called additional 30 DNVs (4.6% increment) (**Table 1, GATK4 + GRAF + DeepTrio + Force-Calling**).

### Sanger Sequencing Confirmations

Ninety-two variants (both FPs and TPs based on visual inspection of the BAM files and thresholds for the second QC step) were chosen for the Sanger sequencing confirmation of the novel DNVS and to investigate the empirical thresholds we developed for the second QC filtering. Because many of the regions harboring these variants were complex, designing successful PCR assays proved technically challenging.

As a result, we were only able to amplify sequences for 73 of our 92 target variants.

*Novel DNVs.* Our three-method workflow revealed 30 novel *de novo* variants, which were not identified by the previous PCGC pipeline (Richter et al., 2020). All these variants were posited as *de novo* based on visual inspection of the BAM files in IGV. Of the 30 novel *de novo* candidates, 23 variants were successfully amplified with PCR. Of those 23, 20 variants were confirmed as *de novo*. Two variants were not found in the proband, and one variant appeared to be inherited (**Table 2, Table S4**).

**Table 2.**
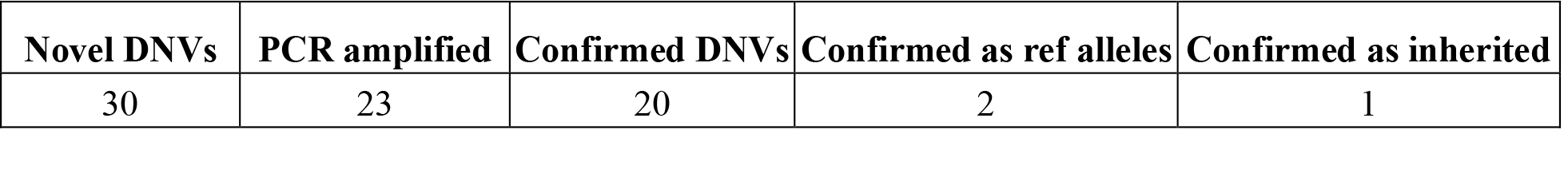
Novel DNV confirmations.

*Alternate alleles in homozygous variants in parents (AAHP).* We assigned upper thresholds for AAC of parents as 1 for SNVs and 0 for indels to remove variants caused by sequencing errors at the second QC filtering step. Twelve variants (both putative FPs and TPs) affected by the threshold were successfully amplified with PCR (**Table S5**). The consistency of the applied thresholds was confirmed for 11 variants.

Three variants yielded inconclusive data for *de novo* filter thresholds, likely due to alignment errors. Two of these variants, initially labeled as inherited, were verified as reference alleles in the proband. One variant, initially marked as inherited from the mother, was confirmed to have been inherited from the father, despite the absence of allele-supporting reads at the location (**Table S5**).

Only one variant, where maternal alignment had the reads with an alternate allele, was found as *de novo*. Therefore, the PCR results showed that our parental AAHP filter was efficient for removing FP variants (**Table 3**).

**Table 3.**
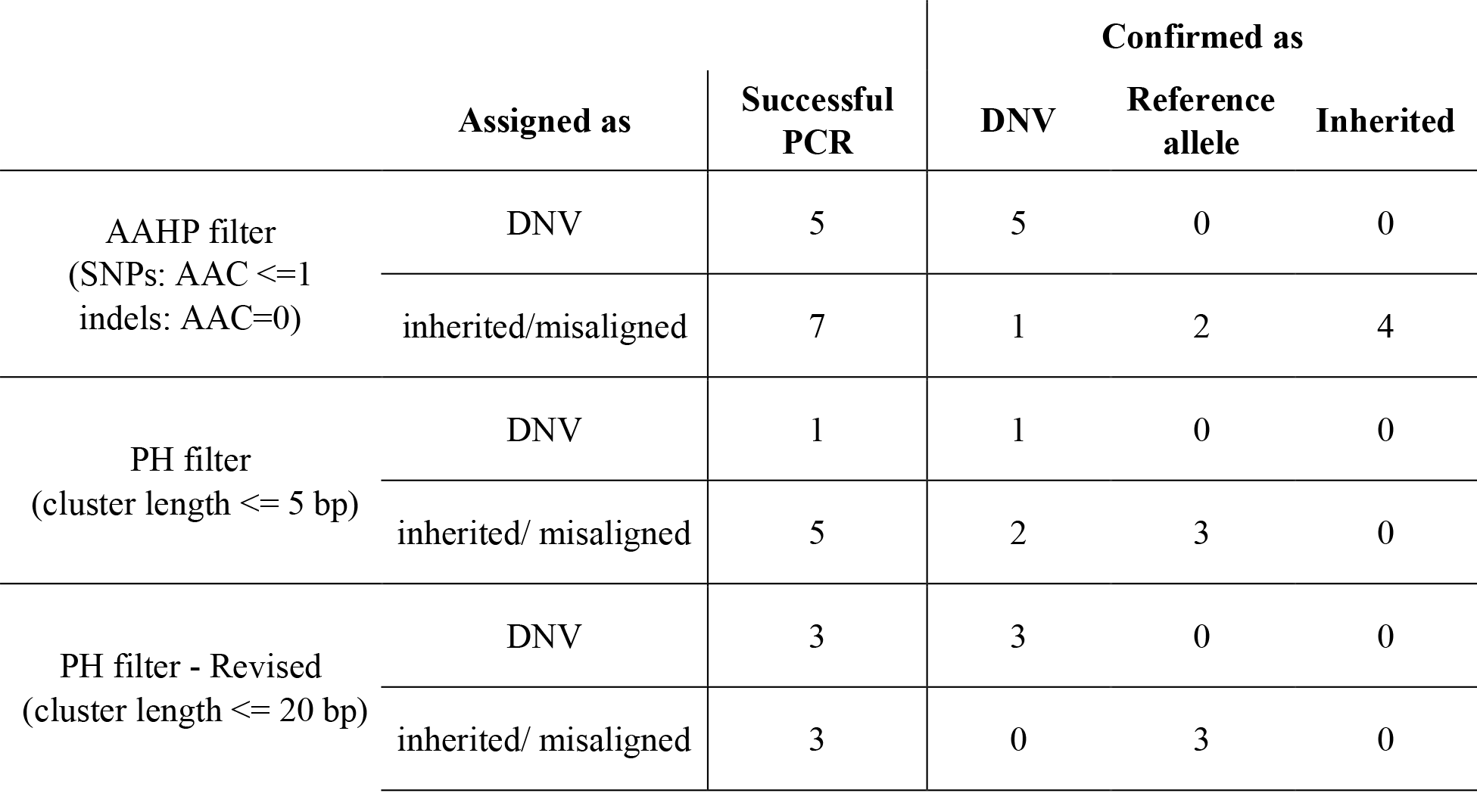
Filter performance.

*Proband haplotype variants.* We considered haplotype variants in probands as FP if the furthest mutations within a haplotype were located > 5 bp apart. Six variants (both FP and TP) affected by the threshold were successfully amplified with PCR (**Table 3, Table S6**). Three variants confirmed as TPs had inter-variant distances of 4, 6 and 11 bp. Another three variants with inter-variant distance of 20, 29 and 31 bp, which we had treated as FP, were confirmed as FPs. The results suggest that the proband haplotype threshold distance might be increased to as much as 12 to 20 bp.

### Overview of *de novo* Variants

We calculated the relative frequencies of mutation classes (**Table 4A, B**), which are in good accordance with mutation spectra previously published (Sasani et al., 2019). Of the 671 DNVs we found, there were 56 variants from UTR and ncRNA regions, 261 intronic DNVs, and 343 intergenic ones (**Table 4C**).

**Table 4.** Statistics of *de novo* variants found in the three-method workflow.

| **A. Mutation class** | **Count** | **Average per sample** | **Relative frequency** |
| --- | --- | --- | --- |
| SNP | 615 | 61.5 | 0.92 |
| SNP coding | 9 | 0.9 | 0.02 |
| indel | 56 | 5.6 | 0.08 |
| indel coding | 0 | 0 | 0 |

| **B. Mutation class** | **Count** | **Average per sample** | **Relative frequency** |
| --- | --- | --- | --- |
| C>T | 166 | 16.6 | 0.25 |
| T>C | 158 | 15.8 | 0.24 |
| CpG>TpG | 75 | 7.5 | 0.11 |
| C>A | 68 | 6.8 | 0.10 |
| C>G | 62 | 6.2 | 0.09 |
| T>G | 47 | 4.7 | 0.07 |
| T>A | 39 | 3.9 | 0.06 |

| **C. Mutation region** | **Count** | **Average per sample** | **Relative frequency** |
| --- | --- | --- | --- |
| exonic | 9 | 0.9 | 0.01 |
| ncRNA | 47 | 4.7 | 0.07 |
| UTR3/UTR5 | 9 | 0.9 | 0.01 |
| intronic | 261 | 26.1 | 0.39 |
| downstream/upstream | 7 | 0.7 | 0.01 |
| intergenic | 343 | 34.3 | 0.51 |

## DISCUSSION

The exploration of DNVs in a proband with any given trait by comparing that person’s genome sequence with those of the unaffected parents is conceptually straightforward. However, identification of less than one hundred DNVs among millions of inherited variants is challenging (Acuna-Hidalgo et al., 2016). Due to sequencing and alignment errors, some variants can be wrongly identified as *de novo*.

Significant efforts have been devoted to improving the efficiency of DNV detection. Earlier methodologies have explored the use of variant annotations, specifically genotype likelihoods, from proband’s and parents’ samples to calculate confidence values for DNVs (Ramu et al., 2013; Wei et al., 2015). While this approach is robust, it often leads to a high number of spurious DNVs, and the final DNV set is dependent on parameters that require tuning by the user. Machine learning techniques show promising results in achieving high accuracy for DNV detection (Khazeeva et al., 2022; Liu et al., 2014). However, the application of machine learning approaches remains challenging due to the insufficient availability of training sets with enough samples. Recently, consensus approaches have emerged with encouraging outcomes (Ng et al., 2022). Our work advances consensus methodologies by incorporating pangenome analysis, introducing a parameter-free automated framework for DNV detection. Such workflow allows processing large numbers of trio GS to call DNVs without needing to undertake visual inspections of the BAM files. We have also designed our methodology to be robust in the presence of mosaicism occasionally observed in the germline tissue samples, by eschewing stringent filtering criteria in favor of force calling because it admits small proportion of alternate alleles in the parents’ samples at putative DNV sites.

Given GS data of a particular average read depth and quality, secondary analyses done by different read aligners and variant callers produce notably different lists of DNVs. In this study, we used a combination of three independent methods to filter out Mendelian violations caused by sequencing or alignment errors.

Only DNVs found by at least two methods were considered further, significantly reducing the number of DNV candidates (the *Consensus* step in **Figure 1**). Although we likely filtered out a small number of TP DNVs, we removed most of FP variants, giving preference to precision over sensitivity. The percentage of the DNVs after the consensus step between the three methods is estimated as 96.7% (**Table 1**). Notably, relaxing thresholds at the first QC step performed individually for each method can increase sensitivity, but the obvious tradeoff is decreased precision. The final step consisted of the re-calling of variant candidates in trios. The second QC filtering (the *force-calling filter* in **Figure 1**) was designed to remove FPs persistent within the three methods, mostly alignment errors. Using any combination of the two methods did not affect precision but notably decreased sensitivity. For the 10 trios studied, the combination of GRAF/DeepTrio revealed 19 fewer DNVs than the three-method workflow, whereas the combinations of GATK4/GRAF and GATK4/DeepTrio missed 49 and 58 DNVs, respectively. Therefore, in the case of limited computational resources, the two-method approach combining the GRAF and DeepTrio pipelines is the most efficient and least expensive (**Table 1**). We expect that an addition of other independent methods (i.e., implementing a workflow with four or more methods) would slowly increase the number of DNVs called but require increasing resources to run. Compared with the previous PCGC study, which used a different approach including convolutional neural network trained on manually curated IGV plots (Richter et al., 2020), our current pipeline identified 99.1% of those DNVs and found 4.6% additional DNVs (**Table 1**).

Applying the three-method workflow, we found 61.5 *de novo* SNPs and 5.6 *de novo* indels per proband, consistent with the expected 44-82 *de novo* SNVs and 3-9 *de novo* indels per individual (Acuna-Hidalgo et al., 2016; Goldmann et al., 2019). Similarly, 0.9 coding *de novo* SNVs per proband were observed, in line with the expected 1-2 per individual (**Table 4A**).

Notably, we focused on only the *de novo* calls where parents had homozygous reference genotypes and proband had heterozygous alternative genotypes because our primary focus is on identifying pathogenic *de novo* mutations. Healthy parents are expected to have reference alleles at the same location where there is a *de novo* variant causing disease in the proband. Therefore, we applied a genotype filter to keep only such *de novo* candidates and determined the quality thresholds accordingly for all callers in the first QC step. We verified that the presence of reads with alternative alleles in parents is a strong indicator of inherited variants, even when the parental genotype is homozygous reference. A variant detected in a related sample acts as a robust prior for a putative variant allele, even when the evidence within the sample itself is limited. This underscores the significance of integrating information from related samples into secondary analysis for *de novo* variant filtering and overall variant calling. We also demonstrated that *de novo* sequence alterations can occur synchronously within regions as wide as 20 base pairs. This represents, to our knowledge, a novel observation in the context of DNVs and provides a crucial insight for future reference.

Coding DNVs are an important cause of Mendelian genetic diseases associated with DNVs as they disrupt or alter gene functions (Deciphering Developmental Disorders Study, 2017; Gilissen et al., 2014; Iossifov et al., 2014). However, they do not explain all the cases. For example, studies of severe, undiagnosed development disorders in children showed that only 42% of individuals carry pathogenic DNVs in coding sequences ^3^. The contribution of non-coding DNVs to the diseases remains to be explored, and we expect that the method presented here will encourage such studies because of its completely automated and scalable processing as well as very high precision and sensitivity of the results.

Overall, the implemented workflow provides a simple and flexible way to investigate DNVs in trios; it retrieves a robust set of DNVs *de novo* from thousands of variant candidates and efficiently filters out Mendelian violations caused by alignment or sequencing errors without requiring manual inspection of variants, thus enabling scalable analysis of large datasets of trio GS.

## ACKNOWLEDGMENTS

We would like to acknowledge Amanda McPartland for her assistance with the Sanger confirmations. PCGC Grants: U01HL153009, 5U01HL128711, U01HL098147.

