## Supplementary_Material for "Automated Identification of Germline *de novo* Mutations in Family Trios: A Consensus-Based Informatic Approach"

Automated *de novo* Mutation Identification

### Content

**Figure S1.** All steps performed in the GATK4, GRAF and DeepTrio pipelines.

**Graph Pangenome Reference**

**Figure S2.** The architecture of graph genome.

**GRAF *de novo* Variant Detection Pipeline**

**Figure S3.** GRAF *de novo* Variant Detection Pipeline

**Figure S4.** CPU Hour distribution for all method combinations

**Table S1.** Tools run in each variant calling pipeline.

**Table S2.** Filtering criteria for quality steps

**Table S3.** Definitions for the metrics used for filtering.

**Table S4.** Sanger sequencing results for the new TP variants were found in addition to the Freeze set.

**Table S5.** Sanger sequencing results for the TP and FP variants affected by the parent AAC threshold.

**Table S6.** PCR results for the TP and FP variants affected by the proband haplotype threshold.

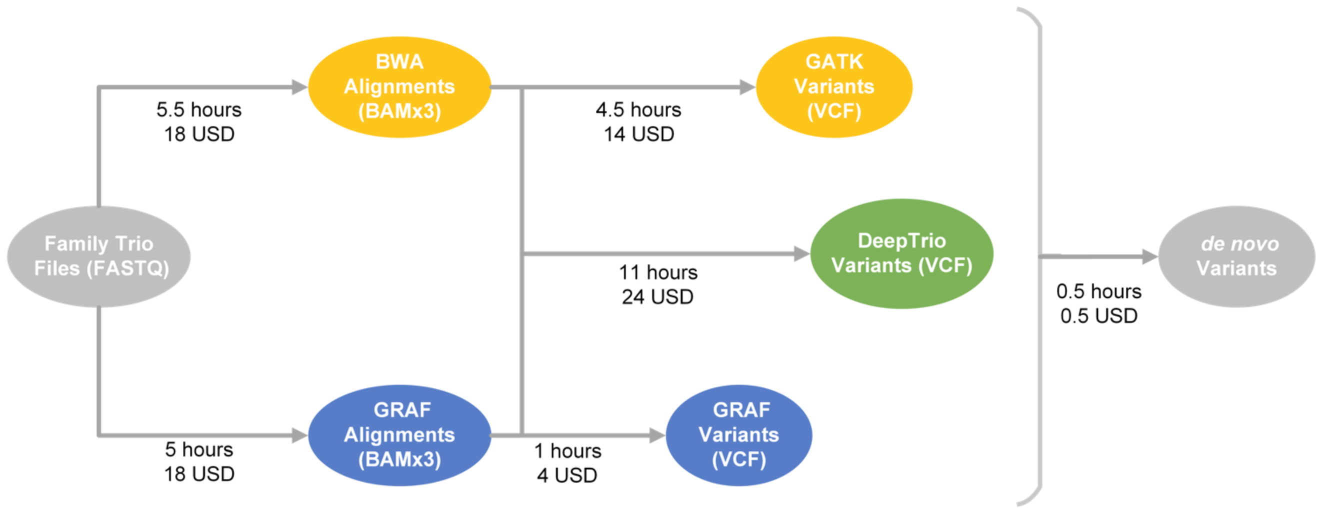


**Figure S1.** All steps performed in the GATK4, GRAF and DeepTrio pipelines. The cost and time were estimated per trio on the Velsera Cavatica platform using spot instances.

#### Graph Pangenome Reference

Standard methods to recover the underlying genome from high throughput short read sequencing samples rely on a single haplotype reference genome assembly to map the sequencing reads to the corresponding generating regions, and then to identify the most likely haplotypes implied by the mapped reads. This process works well for most conserved regions in the genome but fails in regions where the haplotypes in the individual genome are dissimilar to the single haplotypes comprising reference assembly due to evolutionary or somatic divergence. This reference bias, more pronounced in non-European individuals, impairs the identification of relevant genetic mutations for appropriate clinical decision making.

Pangenome references, consisting of multiple haplotypes identified in an appropriately representative cohort of individuals, have been proposed to drastically reduce reference bias inherent in all analyses methods that use reference haplotypes. We have previously described our implementation of pangenome reference-based sequencing data analysis workflow (Rakocevic et al., 2019). We represent the pangenome as a directed graph structure with the chromosomal reference haplotype as the main path in the graph, and the alternate sequences corresponding to genomic variation represented as edges spanning variable genomic regions, as shown in Figure S2. Our representation preserves the genomic loci of genetic variants, ensuring compatibility with existing downstream workflows as well as ease of comparative benchmarking of results with standard methods. The pangenome reference was constructed by augmenting each GRCh38 chromosomal haplotype with associated alt contigs, and with high confidence variants selected from the results reported by 1000 Genomes Phase 3 study (Rakocevic et al., 2019), Simons Genome Diversity Project (Katsnelson, 2010), and other variant datasets (Mallick et al., 2016; Mills et al., 2006).

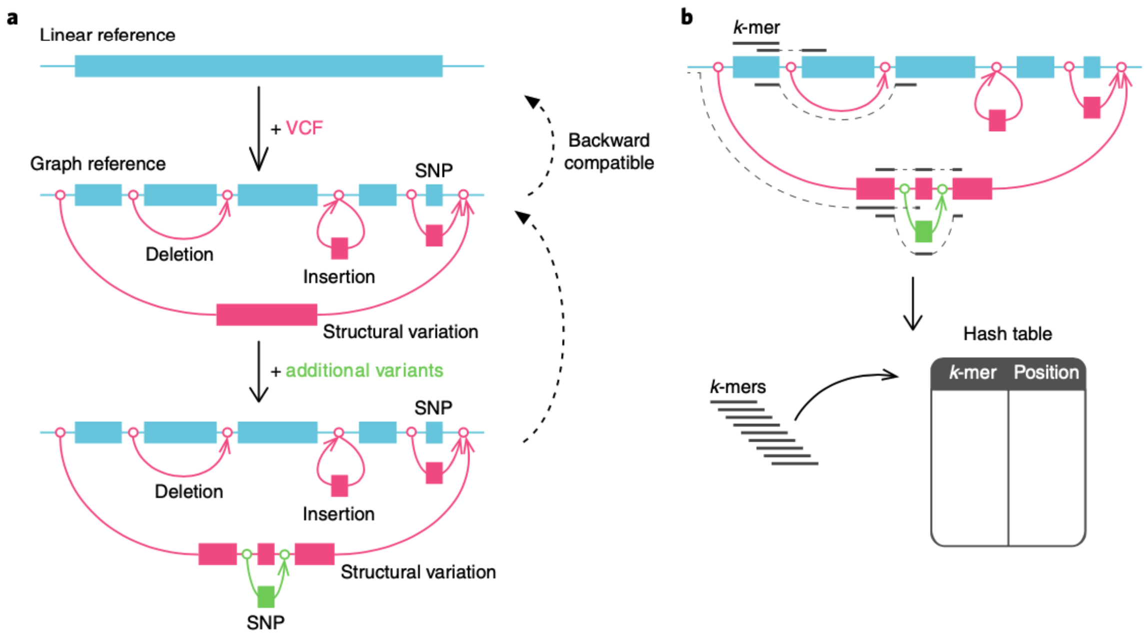


**Figure S2.** The architecture of graph genome. **a.** A set of genetic variants provided in VCF format are added to a standard linear reference genome FASTA file to create a graph genome. Loci on each generated graph genome map unambiguously to corresponding loci on reference haplotype. **b.** Graph genome is indexed by hashes of k-mers along each of the graph's potential pathways mapped against their graph genome loci. Sequencing reads are mapped against the graph by identifying the graph region that shares most k-mers with the read sequence.

#### GRAF *de novo* Variant Detection Pipeline

The steps for filtering *de novo* variants for the GRAF pipeline are as follows:

1. Merge family VCFs into a multi-sample family VCF file. We used Bcftools Merge (Danecek et al., 2021) for this task (Merge Variants)
2. Identify the de novo variants within the family VCF file:

2.1. For each variant in the family VCF file, compare the child genotype with each parent's genotype respectively. We used RTG VCFeval (Cleary et al., 2015) tool for this task (Compare Variants)

2.2. If the child genotype differs from either parent's genotype, inspect each genotype triple comprising the variant call for Mendelian inconsistency to get a list of putative de novo variant loci (Collect Mendelian Violation (MV) Loci)

2.3. Use the candidate de novo variant loci to filter family VCF (Filter by Region)

2.4. Collect Mendelian violations from the family VCF filtered with candidate de novo loci. We use RTG Mendelian tool for this task (Collect Mendelian Violations (MVs))

2.5. Further reduce this list by eliminating all genotype triples where any of 6 alleles comprising the 3 sample genotypes does not have an adequate proportion of mapped reads containing evidence for the allele (Filter by Allelic Balance (AB) and Mapped Allelic Balance (MAB))

Figure S3 shows the steps for filtering *de novo* variants for GRAF pipeline.

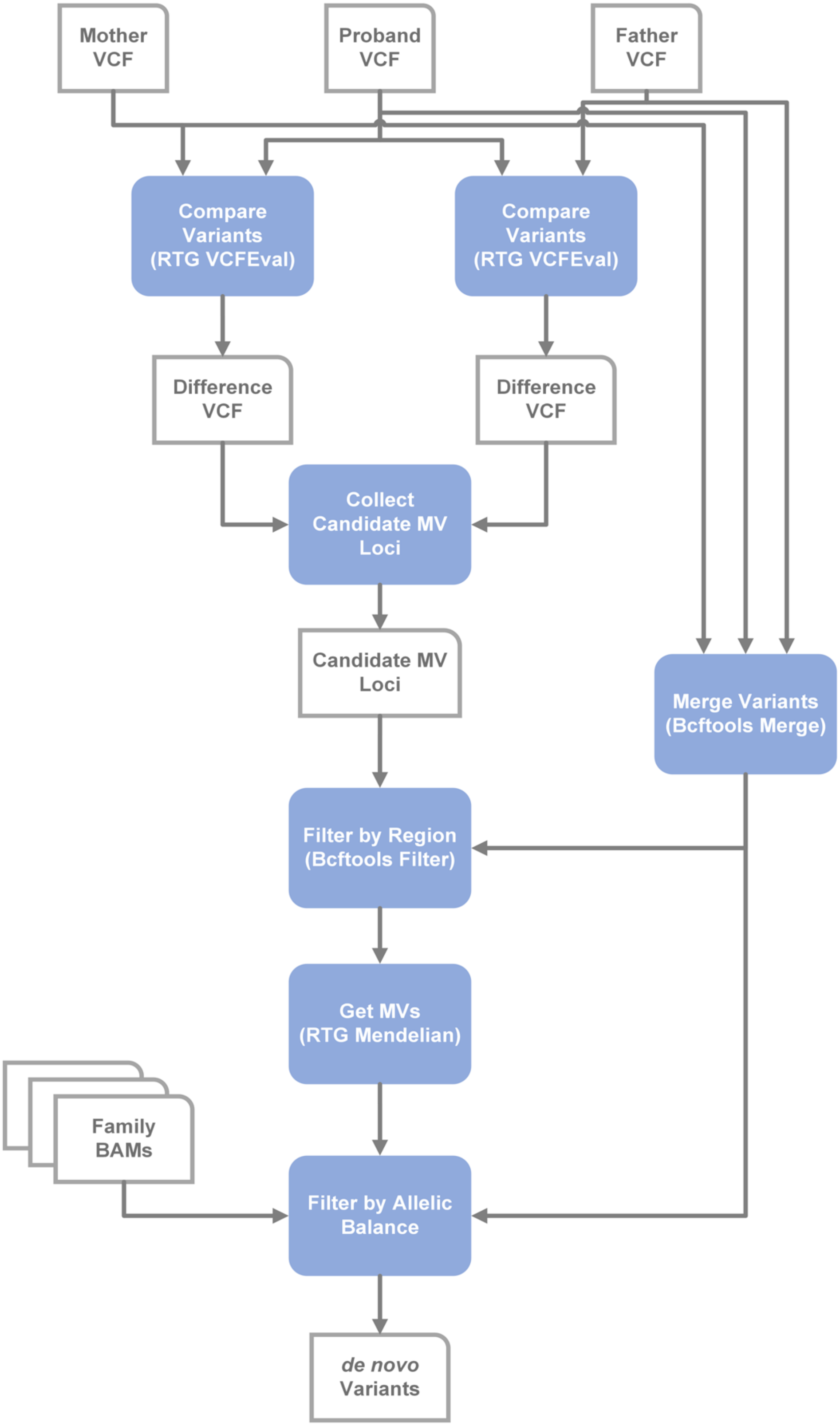


**Figure S3.** GRAF *de novo* Variant Detection Pipeline

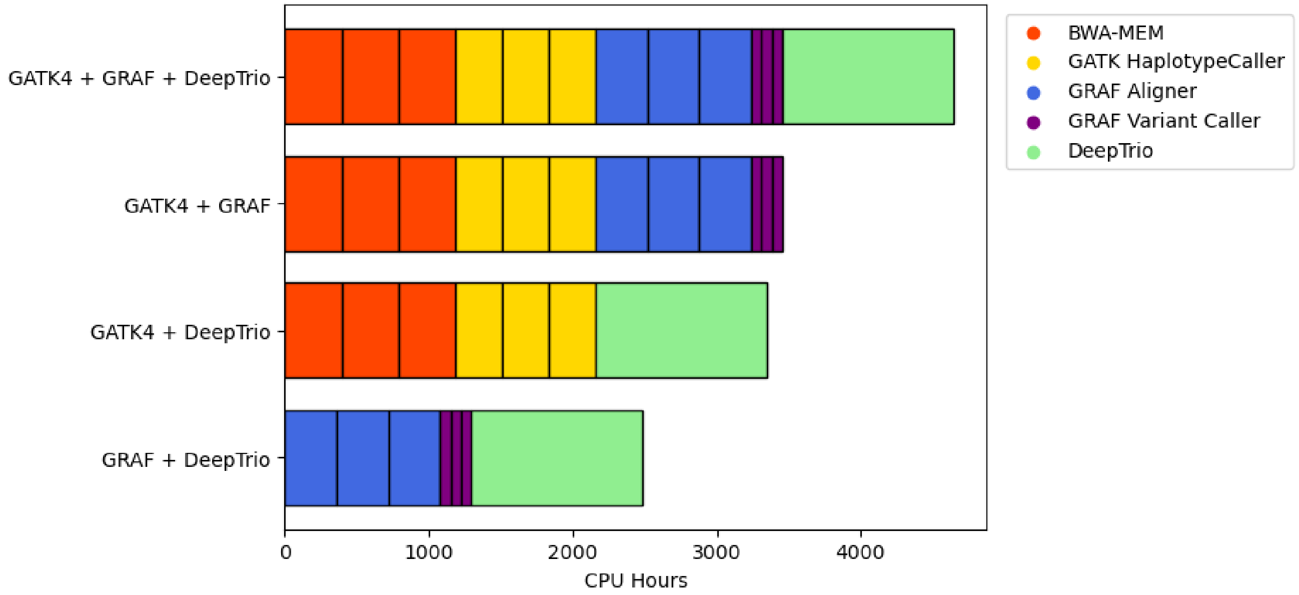


**Figure S4.** CPU Hour distribution for all method combinations

**Table S1.** Tools run in each variant calling pipeline.

| | **Alignments** | **Variant Calling** |
|---|---|---|
| **GATK4** | PICARD FastqToSam<br>PICARD MarkIlluminaAdapters<br>PICARD SamToFastq<br>BWA-MEM<br>GATK MergeBamAlignment<br>GATK MarkDuplicates<br>GATK SortSam<br>GATK SetNmMdAndUqTags<br>GATK BaseRecalibrator<br>GATK ApplyBQSR | GATK HaplotypeCaller<br>GATK GenomicsDBImport<br>GATK GenotypeGVCFs<br>GATK VariantRecalibrator<br>GATK ApplyVQSR<br>GATK CalculateGenotypePosteriors<br>GATK VariantFiltration<br>GATK VariantAnnotator<br>BCFTools Norm<br>Quality Filter |
| **GRAF** | GRAF Aligner | GRAF VariantCaller<br>GRAF *de novo* Detection Pipeline |
| **DeepTrio** | PICARD FastqToSam<br>PICARD MarkIlluminaAdapters<br>PICARD SamToFastq<br>BWA-MEM<br>GATK MergeBamAlignment<br>GATK MarkDuplicates<br>GATK SortSam<br>GATK SetNmMdAndUqTags<br>GATK BaseRecalibrator<br>GATK ApplyBQSR | DeepTrio<br>GLNexus<br>BCFTools Norm<br>Quality Filter |

**Table S2.** Filtering criteria for quality steps

| | | First QC | | | Second QC |
|---|---|---|---|---|---|
| **Filter name** | **Definition** | **GATK4** | **GRAF** | **DeepTrio** | **Force-calling** |
| *de novo* candidate | Genotype criteria | Biallelic: Parents (0/0), Proband (0/1) | Biallelic: Parents (0/0), Proband (0/1) | Biallelic: Parents (0/0), Proband (0/1) | Biallelic: Parents (0/0), Proband (0/1) |
| *de novo* candidate in chrX for male probands | Genotype criteria | Monoallelic: Parents (0), Proband (1) | Monoallelic: Parents (0), Proband (1) | Monoallelic: Parents (0), Proband (1) | Monoallelic: Parents (0), Proband (1) |
| Variant filters | Additional filter parameters | PASS, hiConfDeNovo loConfDeNovo | - | --config DeepVariant_unfiltered | Cluster length in proband ≤ 5 bp |
| AC | Allele Count | 1 | - | 1 | 1 |
| QD | Quality by depth | - | - | - | - |
| Parents GQ | Genotype quality | - | - | - | - |
| Proband GQ | Genotype quality | ≥ 60 | - | ≥ 0 | ≥ 60 |
| Parents DP | Variant read depth | ≥ 7 | - | ≥ 7 | ≥ 5 |
| Proband DP | Variant read depth | ≥ 7 | - | ≥ 7 | ≥ 10 |
| Parents AB | Allelic balance | ≤ 0.01 | - | ≤ 0.1 | ≤ 0.15 |
| Proband AB | Allelic balance | $0.2 \leq x \leq 0.8$ | $\forall x \geq 0.15$ & $\Sigma x \geq 0.9$ | $0.2 \leq x \leq 0.8$ | $0.2 \leq x \leq 0.8$ |
| Parents AAD | Alternative Allele Depth | - | - | - | ≤ 1 for SNVs =0 for indels |
| Proband AAD | Alternative Allele Depth | ≥ 7 | - | ≥ 7 | ≥ 5 |

| | | | | | |
|---|---|---|---|---|---|
| Parents MAB | Mapped Allelic Balance | | $\Sigma x \geq 0.9$ | | |
| Proband MAB | Mapped Allelic Balance | - | $\forall x \geq 0.15$ & $\Sigma x \geq 0.9$ | - | - |

**Table S3.** Definitions for the metrics used for filtering.

| **Metric Name** | **Definition** |
|---|---|
| AC | Allele Count |
| QD | Quality by depth, variant call quality normalized by read depth |
| GQ | Variant's genotype quality |
| DP | Read depth of the variant, total number of reads aligned to variant location |
| AB | Allelic balance for the variant, ratio of alt supporting reads to the total depth in the VCF |
| AAD | Alternate allele depth, number of reads supporting the variant in the VCF |
| MAB | Number of reads supporting the alleles in the read pile-up to the total number of reads |

**Table S4.** Sanger sequencing results for the new TP variants were found in addition to the Freeze set.

| **Variant** | | | | | **Computational results** | | | | | **PCR results** | |
|---|---|---|---|---|---|---|---|---|---|---|---|
| **Chrom** | **Pos** | **Ref** | **Alt** | **Family** | **Pipelines** | **AD$^{pat}$ * (AB)** | **AD$^{mat}$ * (AB)** | **AD$^{Pro}$ * (AB)** | **Prediction** | **Proband** | ***de novo*** |
| chr2 | 42,384,880 | G | T | 1-04537 | GATK4, GRAF | 38/0 (0) | 27/0 (0) | 27/7 (0.21) | *de novo* | Reference allele | Not correct |
| chr2 | 203,431,709 | C | A | 1-04537 | GATK4, GRAF, DeepTrio | 40/0 (0) | 40/0 (0) | 18/15 (0.45) | *de novo* | Inherited (pat) | Not correct |
| chr3 | 90,631,023 | G | A | 1-04460 | GATK4, GRAF, DeepTrio | 60/0 (0) | 41/0 (0) | 24/26 (0.52) | *de novo* | Reference allele | Not correct |
| chr3 | 91,458,964 | G | T | 1-00801 | GATK4, GRAF, DeepTrio | 42/0 (0) | 42/0 (0) | 21/28 (0.57) | *de novo* | *de novo* | Confirmed |
| chr6 | 85,868,792 | C | T | 1-01019 | GATK4, GRAF, DeepTrio | 32/0 (0) | 40/0 (0) | 17/20 (0.54) | *de novo* | *de novo* | Confirmed |
| chr7 | 131,070,605 | T | C | 1-04460 | GATK4, GRAF, DeepTrio | 28/0 (0) | 43/0 (0) | 19/12 (0.39) | *de novo* | *de novo* | Confirmed |
| chr8 | 95,414,588 | CCATT | C | 1-04389 | GATK4, GRAF, DeepTrio | 37/0 (0) | 34/0 (0) | 18/18 (0.5) | *de novo* | *de novo* | Confirmed |
| chr8 | 105,748,089 | T | C | 1-04537 | GATK4, GRAF, DeepTrio | 31/0 (0) | 51/0 (0) | 16/17 (0.52) | *de novo* | *de novo* | Confirmed |
| chr9 | 28,661,917 | G | A | 1-04389 | GATK4, GRAF, DeepTrio | 46/0 (0) | 40/0 (0) | 16/16 (0.5) | *de novo* | *de novo* | Confirmed |
| chr10 | 33,144,531 | ATTAG | A | 1-04389 | GATK4, GRAF, DeepTrio | 36/0 (0) | 41/0 (0) | 18/21 (0.54) | *de novo* | *de novo* | Confirmed |
| chr10 | 124,132,804 | T | G | 1-04190 | GATK4, GRAF, DeepTrio | 27/0 (0) | 27/0 (0) | 18/21 (0.54) | *de novo* | *de novo* | Confirmed |
| chr14 | 43,735,998 | G | C | 1-03897 | GRAF, DeepTrio | 10/0 (0) | 28/0 (0) | 8/15 (0.65) | *de novo* | *de novo* | Confirmed |
| chr17 | 63,343,374 | A | G | 1-05846 | GATK4, GRAF, DeepTrio | 35/0 (0) | 34/0 (0) | 16/18 (0.53) | *de novo* | *de novo* | Confirmed |

| | | | | | | | | | | | |
|---|---|---|---|---|---|---|---|---|---|---|---|
| chr18 | 45,216,791 | C | T | 1-05673 | GATK4, GRAF, DeepTrio | 41/0 (0) | 36/0 (0) | 23/18 (0.44) | *de novo* | *de novo* | Confirmed |
| chr18 | 72,806,285 | C | T | 1-05443 | GATK4, GRAF | 39/0 (0) | 49/0 (0) | 32/8 (0.2) | *de novo* | *de novo* (mosaic) | Confirmed |
| chr19 | 17,896,000 | C | A | 1-00801 | GRAF, DeepTrio | 31/0 (0) | 16/0 (0) | 14/12 (0.46) | *de novo* | *de novo* | Confirmed |
| chrX | 3,712,869 | C | T | 1-04190 | GRAF, DeepTrio | 12/0 (0) | 33/0 (0) | 22/12 (0.35) | *de novo* | *de novo* | Confirmed |
| chrX | 5,450,254 | C | T | 1-05443 | GRAF, DeepTrio | 15/0 (0) | 50/0 (0) | 25/18 (0.42) | *de novo* | *de novo* | Confirmed |
| chrX | 29,087,338 | A | G | 1-05846 | GRAF, DeepTrio | 18/0 (0) | 34/0 (0) | 23/16 (0.41) | *de novo* | *de novo* | Confirmed |
| chrX | 51,565,010 | G | GT | 1-04190 | GRAF, DeepTrio | 18/0 (0) | 39/0 (0) | 12/21 (0.64) | *de novo* | *de novo* | Confirmed |
| chrX | 83,704,567 | G | A | 1-04190 | GRAF, DeepTrio | 17/0 (0) | 32/0 (0) | 16/25 (0.61) | *de novo* | *de novo* | Confirmed |
| chrX | 126,272,355 | C | A | 1-01019 | GRAF, DeepTrio | 19/0 (0) | 38/0 (0) | 27/20 (0.43) | *de novo* | *de novo* | Confirmed |
| chrX | 129,316,052 | A | G | 1-05443 | GRAF, DeepTrio | 14/0 (0) | 35/0 (0) | 30/19 (0.39) | *de novo* | *de novo* | Confirmed |

* *AD = ($AD^{ref}$/ $AD^{alt}$)*

**Table S5.** Sanger sequencing results for the TP and FP variants affected by the parent AAC threshold.

| **Variant** | | | | | **Computational results** | | | | | **PCR results** | |
|---|---|---|---|---|---|---|---|---|---|---|---|
| **Chrom** | **Pos** | **Ref** | **Alt** | **Family** | **Pipelines** | **AD$^{pat}$ * (AB)** | **AD$^{mat}$ * (AB)** | **AD$^{Pro}$ * (AB)** | **Prediction** | **Result** | **Threshold **** |
| chr4 | 86,330,549 | G | C | 1-04190 | Freeze, GATK4, GRAF, DeepTrio | 23/0 (0) | 38/1 (0.03) | 20/25 (0.56) | *de novo* | *de novo* | Confirmed |
| chr5 | 31,218,501 | C | G | 1-03897 | Freeze, GATK4, GRAF, DeepTrio | 31/0 (0) | 50/1 (0.02) | 23/15 (0.39) | *de novo* | *de novo* | Confirmed |
| chr9 | 107,787,103 | C | CTTTTTT | 1-05443 | GRAF, DeepTrio | 8/0 (0) | 2/1 (0.33) | 7/1 (0.13) | Inherited (mat) | Inherited (mat) | Confirmed |
| chr11 | 95,026,363 | T | G | 1-04537 | Freeze, GATK4, GRAF, DeepTrio | 33/0 (0) | 36/1 (0.03) | 27/15 (0.36) | *de novo* | *de novo* (mosaic) | Confirmed |
| chr12 | 33,275,328 | C | CTTTTTTTTT | 1-04460 | GATK4, GRAF, DeepTrio | 14/3 (0.18) | 21/0 (0) | 4/10 (0.71) | Inherited (pat) | Inherited (pat) | Confirmed |
| chr13 | 44,111,968 | T | G | 1-04537 | Freeze, GATK4, GRAF, DeepTrio | 40/0 (0) | 44/1 (0.02) | 31/10 (0.24) | *de novo* | *de novo* | Confirmed |
| chr13 | 63,468,666 | C | T | 1-04389 | Freeze, GATK4, GRAF, DeepTrio | 36/1 (0.03) | 33/0 (0) | 20/27 (0.57) | *de novo* | *de novo* | Confirmed |
| chr13 | 112,164,593 | T | C | 1-05443 | GATK4, GRAF | 25/2 (0.07) | 29/1 (0.03) | 28/10 (0.26) | Inherited (pat,mat) | Reference allele | Alignment error |
| chr14 | 38,185,150 | T | C | 1-04460 | GATK4, GRAF, DeepTrio | 31/0 (0) | 31/3 (0.09) | 22/14 (0.39) | Inherited (mat) | *de novo* | Not correct |
| chr19 | 57,319,997 | C | CTTTTTTTTTTT | 1-03897 | GATK4, GRAF | 18/0 (0) | 15/2 (0.12) | 11/7 (0.39) | Inherited (mat) | Inherited (pat,mat) | Partially confirmed |
| chr20 | 35,881,048 | C | CTTTTTTT | 1-05846 | GRAF, DeepTrio | 14/0 (0) | 7/2 (0.22) | 4/4 (0.5) | Inherited (mat) | Reference allele | Alignment error |
| chr20 | 42,299,641 | C | CTTTT | 1-01019 | GRAF, DeepTrio | 10/0 (0) | 5/2 (0.29) | 7/5 (0.42) | Inherited (mat) | Inherited (pat) | Alignment error |

** AD = (AD$^{ref}$/ AD$^{alt}$)*

*** AAC of parents <= 1 for SNVs and 0 for indels*

**Table S6.** PCR results for the TP and FP variants affected by the proband haplotype threshold.

| Variant | | | | | Computational results | | | | PCR results | |
|---|---|---|---|---|---|---|---|---|---|---|
| **Chrom** | **Pos** | **Ref** | **Alt** | **Family** | **Pipelines** | **# of events** | **Length** | **Prediction** | **Result** | **Threshold*** |
| chr1 | 14,251,90; 14,251,91 | C; C | T; G | 1-04389 | Freeze, GRAF, DeepTrio; GATK4, GRAF, DeepTrio | 3 | 11 b3p | Alignment Error | *de novo* | Not correct |
| chr2 | 148,822,048 | A | C | 1-00801 | GATK4, GRAF | 7 | 20 bp | Alignment Error | Reference allele | Confirmed |
| chr8 | 110,371,875; 110,371,878 | G; G | A; A | 1-00801 | Freeze, GATK4, GRAF, DeepTrio; Freeze, GATK4, GRAF, DeepTrio | 2 | 4 bp | multiple *de novo* mutations | *de novo* | Confirmed |
| chr12 | 50171621; 50171626 | T; A | C; G | 1-04389 | Freeze, GRAF; Freeze, GRAF | 5 | 29 bp | Alignment Error | Reference allele | Confirmed |
| chr14 | 89,040,462; 89,040,466; 89,040,468 | A C; A; C | A; T; G | 1-05673 | GATK4, GRAF, DeepTrio; Freeze, GATK4, GRAF, DeepTrio; Freeze, GATK4, GRAF, DeepTrio | 3 | 6 bp | Alignment Error | *de novo* | Not correct |
| chrX | 1,096,471 | C | A | 1-04460 | GATK4, GRAF | 4 | 31 bp | Alignment Error | Reference allele | Confirmed |

** Haplotype variants in probands are considered as FP if the furthest mutations within a haplotype were located > 5 bp apart*
